# Redundant roles of YES and SRC tyrosine kinases in driving malignant peripheral nerve sheath tumors

**DOI:** 10.64898/2026.02.17.706384

**Authors:** Elamine Zereg, Laure Voisin, Mathieu Courcelles, Sylvie Brochu, Melania Gombos, Éric Bonneil, Karl Grenier, Sungmi Jung, Claude Perreault, Franck Tirode, Pierre Thibault, Sylvain Meloche

**Author notes:** **Correspondence**: Dr Sylvain Meloche, Institute for Research in Immunology and Cancer 2950, Chemin de Polytechnique, Montreal, QC H3C 3J7, Canada. The authors declare no competing interests.

## Abstract

Malignant peripheral nerve sheath tumors (MPNST) are highly aggressive soft tissue sarcomas that are largely incurable with no clinically effective systemic therapies or immunotherapies for advanced disease. Here, we identify the SRC-family kinases (SFKs) YES and SRC as redundant, essential drivers of MPNST growth. Dual inhibition of YES/SRC activity by genetic silencing or pharmacological SFK inhibitors markedly suppressed the proliferation of multiple NF1-mutant MPNST cell lines. In vivo, conditional genetic depletion of YES/SRC in MPNST cells abrogated tumor growth in subcutaneous and orthotopic models, and dasatinib treatment delayed tumor progression and improved overall survival. Integrated transcriptomic and phosphotyrosine proteomic analyses revealed that YES/SRC inactivation extensively rewires MPNST signaling, coordinately repressing multiple oncogenic signaling pathways and downstream cell cycle transcriptional programs. Unexpectedly, YES/SRC inhibition also upregulated interferon and antigen processing and presentation pathways and increased cell-surface MHC class I expression, consistent with tumor-intrinsic immune reactivation. Clinically, analysis of a large sarcoma cohort demonstrated that *YES1* is significantly overexpressed in MPNST compared to benign soft tissue tumors. Collectively, our findings establish YES/SRC as non-oncogene vulnerabilities in MPNST.

## INTRODUCTION

Malignant peripheral nerve sheath tumors (MPNSTs) are rare but highly aggressive soft tissue sarcomas that arise from Schwann cell precursors (Yao et al, 2023). Approximately half of all MPNST cases occur in the context of neurofibromatosis type 1 (NF1), while the remainder develop sporadically or following prior radiation exposure. Surgical resection remains the mainstay of treatment, yet recurrence and metastasis are common, leading to poor clinical outcomes and a 5-year overall survival rate of ∼ 50% (Cai et al, 2020; Martin et al, 2020; Yao et al., 2023). Treatment with radiotherapy or chemotherapy is largely ineffective for MPNST patients. MPNSTs are also described as immune-cold tumors with low T-cell infiltration and downregulation of antigen processing and presentation genes, thereby compromising the potential use of immune checkpoint inhibitors (Lee et al, 2006; Paudel et al, 2023).

Comprehensive genomic and transcriptomic analyses have revealed a complex landscape of oncogenic drivers and tumor suppressor losses in MPNSTs (Kim & Pratilas, 2018; Suppiah et al, 2023; Yao et al., 2023). Key tumor suppressor genes frequently inactivated include *NF1*, *CDKN2A/B*, *TP53*, and *PTEN*, while various receptor tyrosine kinases such as epidermal growth factor receptor and platelet-derived growth factor receptor are often activated, and occasionally amplified, in these tumors. These genetic alterations are accompanied by aberrant activation of intracellular signaling pathways such as RAS-ERK1/2 MAP kinase (MAPK), phosphatidylinositol 3-kinase (PI3K)/AKT, mTOR, Wnt/β-catenin, and sonic hedgehog. Additionally, epigenetic dysregulation plays an important role in MPNST progression. Loss-of-function mutations in core components of the polycomb repressive complex 2 (PRC2), notably SUZ12 and EED, are found in a majority of MPNSTs, leading to a global reduction of H3K27me3-mediated transcriptional repression (Lee et al, 2014). Despite significant progress in unraveling the molecular mechanisms underlying MPNST pathogenesis, these findings remain to be translated into clinical practice. There is a pressing need to define actionable vulnerabilities in MPNSTs to guide the development of targeted and immune-based therapies.

More recently, dysregulation of the Hippo-YAP/TAZ pathway has emerged as a driver of Schwann cell transformation. The Hippo signaling pathway is a conserved kinase cascade that negatively regulates the transcriptional coactivators YAP and TAZ, which play a vital role in cell proliferation, differentiation, stem cell self-renewal, and organ homeostasis (Ma et al, 2019; Zheng & Pan, 2019). Human MPNSTs exhibit elevated YAP/TAZ nuclear expression and transcriptional activity (Isfort et al, 2019; Wu et al, 2018). Hyperactivation of YAP/TAZ in Schwann cells, consequent to inactivation of Lats1/2 or Taok1 kinases, can transform Schwann cells and induce high-grade MPNSTs in mice (Velez-Reyes et al, 2021; Wu et al., 2018). Inhibition of YAP/TAZ activity inhibits MPNST cell proliferation and reduces tumor growth. These findings suggest that the Hippo-YAP/TAZ pathway may have therapeutic potential in MPNSTs.

YAP/TAZ factors are regulated by multiple oncogenic signaling pathways that control their nucleocytoplasmic shuttling and the transcription of target genes (Piccolo et al, 2023). Notably, the SRC-family kinases (SFKs) YES and SRC promote the nuclear accumulation and transcriptional activity of YAP/TAZ, either through direct phosphorylation or by repressing LATS1/2 activity (Guegan et al, 2022; Lamar et al, 2019; Li et al, 2016; Rosenbluh et al, 2012; Si et al, 2017). In this study, we investigated the functional relevance of YES and SRC tyrosine kinases in the pathogenesis of MPNST. We found that YES and SRC function redundantly to sustain MPNST cell proliferation and tumor growth. Dual inhibition of YES/SRC signaling downregulated oncogenic signaling pathways and cell division, while upregulating inflammatory signaling and MHC class I expression in MPNST cells. Our findings identify YES/SRC as candidate therapeutic targets for the treatment of MPNST.

## RESULTS

### Depletion of YAP/TAZ impairs the proliferation of plexiform neurofibroma and MPNST cells

Previous work has shown that genetic reduction of YAP/TAZ expression inhibits tumor progression in Lats1/2-deficient nerves, and that treatment with the non-selective YAP-TEAD complex inhibitor verteporfin partially reduces MPNST cell proliferation (Wu et al., 2018). To assess the generalizability of these findings, we examined the effect of siRNAs simultaneously targeting YAP and TAZ on the proliferation of a panel of plexiform neurofibroma and MPNST cell lines. Partial depletion of YAP and TAZ consistently suppressed proliferation in all tumor cell lines tested (Fig. EV1), reinforcing the critical role of YAP/TAZ signaling in sustaining the viability of both benign and malignant nerve sheath tumors. To evaluate the clinical relevance of these findings, we analyzed by immunohistochemistry (IHC) a clinically annotated tissue microarray (TMA) comprising 14 low-grade and 27 high-grade cases of MPNSTs, and 24 benign neurofibroma and schwannoma samples. Using a validated antibody against YAP, or against both YAP and TAZ, we observed a significant increase in YAP/TAZ abundance in MPNSTs compared to benign tumors (Fig. EV2), consistent with previous observations (Wu et al., 2018).

### Pharmacological or genetic inhibition of YES/SRC signaling impairs MPNST cell proliferation

Given the known epistatic interaction between the SFKs YES and SRC and YAP/TAZ signaling, we next asked whether YES and SRC also regulate the proliferation of MPNST cell lines. Treatment with the non-selective SFK inhibitor dasatinib (Kantarjian et al, 2006) inhibited the proliferation of all three human MPNST cell lines tested, with IC_50_ values ranging from 9 to 74 nM (Fig. 1A). Similar anti-proliferative effects were observed with ponatinib, a potent and structurally distinct inhibitor of YES and SRC (O’Hare et al, 2009), suggesting that the effect is on-target (Fig. EV3).

**Figure 1.**
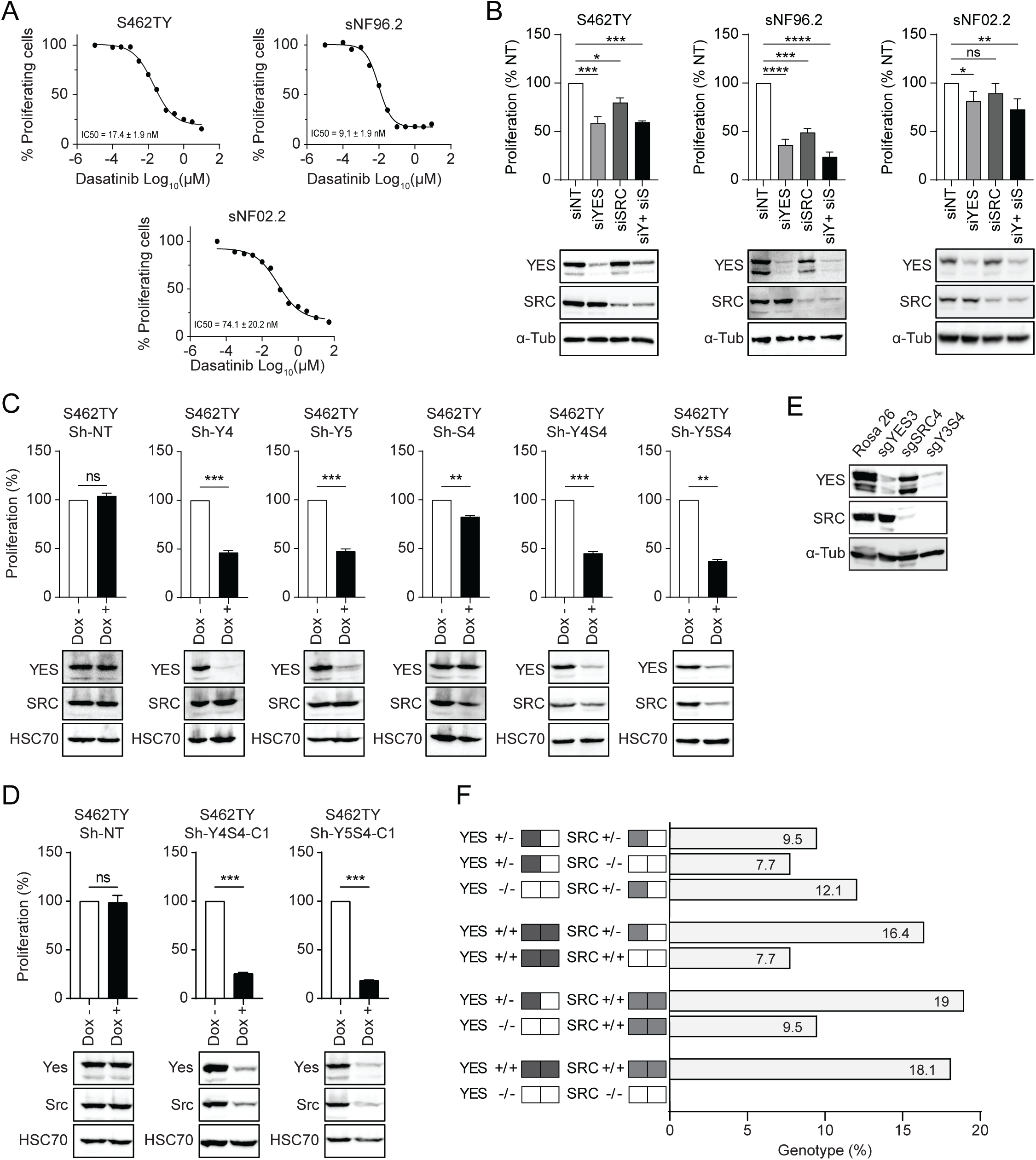
Pharmacological or genetic inhibition of YES/SRC signaling inhibits the proliferation of human MPNST cells. (**A**) Dose-response curves of dasatinib on the proliferation of human MPNST cell lines. IC_50_ values are indicated and represent the mean ± SEM of three experiments. (**B**) MPNST cell lines were transfected with non-target (NT), *YES1*, *SRC*, or combined *YES1*/*SRC* (YS) targeted siRNAs. Cell proliferation was assayed after 5 days. Top, cell proliferation expressed as mean ± SEM relative to NT control (n=3 or 4, one-way ANOVA with Dunnett’s post hoc test). Bottom, representative western blots. (**C**) S462TY cells were infected with lentiviruses expressing non-target control (shNT), *YES1* (Y), *SRC* (S), or combined *YES1*/*SRC* (YS) Dox-inducible shRNAs. Populations of stably infected cells were selected with puromycin. After 5 days, the cells were treated or not with 400 ng/ml doxycycline. Cell proliferation was assayed after 7 days. Top, cell proliferation expressed as mean ± SEM relative to NT control (n=3, paired Student’s t-test). Bottom, representative western blots. (**D**) Cell proliferation assay of two clonal lines of S462TY cells expressing *YES1*/*SRC* Dox-inducible shRNAs treated as in C. (**E** and **F**) S462TY cells were infected with lentiviruses encoding Cas9 and sgRNAs targeting control Rosa26 locus or *YES1*, *SRC*, or combined *YES1*/*SRC* genes. Edited cell populations were selected with puromycin for 4 days. (**E**) Western blot analysis of YES and SRC expression. (**F**) Clonal analysis of cell proliferation. Single-cell derived clones were isolated by limiting dilution and plated in 96-well plates. After 2-3 weeks of culture, 136 cell clones were lysed and analyzed by dot blotting for expression of YES and SRC. Results are expressed as percentage of the total number of clones. ns, not significant. \**P* < 0.05, ** *P* < 0.01, *** *P* < 0.001, **** *P* < 0.0001.

The essential role of YES and SRC was further validated using robust genetic approaches. Transient depletion of YES expression with SMARTpool siRNAs significantly impaired the proliferation of S462TY, sNF96.2 and sNF02.2 MPNST cell lines, while knockdown of SRC produced a more modest effect (Fig. 1B). We then focused our analyses on the S462TY cell line, which shows high sensitivity to SFK inhibitors and reliably form tumors in immunodeficient mice, providing a suitable model for in vivo studies. To confirm the siRNA results, we engineered populations of S462TY cells with validated Dox-inducible shRNAs targeting *YES1*, *SRC* or both genes. YES and SRC proteins have long half-lives and time-course experiments revealed that at least 72 h of doxycycline treatment was required to achieve efficient knockdown of the two kinases (Fig. EV4). Consistent with the effect of siRNAs, conditional depletion of YES with two distinct non-overlapping shRNAs markedly reduced the proliferation of S462TY cells, whereas SRC depletion caused a weaker inhibitory effect (Fig. 1C). To reduce clonal heterogeneity, we isolated two individual clones from S462TY cells co-expressing *YES1* and *SRC*-targeting shRNAs. Despite incomplete silencing of YES and SRC expression, the proliferation of both cell clones was markedly inhibited upon doxycycline treatment (Fig. 1D).

### YES and SRC act redundantly to sustain MPNST cell viability

Combined depletion of YES and SRC by RNAi consistently showed a more robust and reproducible inhibition of MPNST cell proliferation compared to individual kinase targeting. To rigorously assess the functional redundancy of YES and SRC, we co-infected S462TY cells with lentiviruses encoding Cas9 and sgRNAs targeting *YES1* and *SRC*. Western blot analysis confirmed the efficient depletion of both proteins in the edited S462TY cell population (Fig. 1E). To directly test for functional redundancy, single-cell clones were isolated from the edited S462TYcell population by limiting dilution and analyzed for expression of YES and SRC by semi-quantitative dot blotting (Fig. EV5). From the analysis of 136 clones, none of the clones exhibited a complete loss of both *YES1* and *SRC* alleles after expansion (Fig. 1F). These results provide strong genetic evidence for the functional redundancy and essential role of YES and SRC in sustaining the viability of MPNST cells.

### Dual targeting of YES/SRC inhibits tumor growth in subcutaneous and orthotopic MPNST models

To assess the in vivo relevance of these observations, we inoculated NSG mice with two individual clones (Y4S4-C1 and Y5S4-C1) of S462TY cells expressing doxycycline-inducible shRNAs targeting *YES1* and *SRC* (see Fig. 1D). S462TY cells expressing a non-target shRNA were used as control. The cells were injected subcutaneously and once tumors reached a volume of ∼150 mm^3^, the mice were treated with vehicle or doxycycline. Co-depletion of YES and SRC completely abolished tumor growth in both S462TY clones (Fig. 2A). Only a weak effect of doxycycline was observed in cells expressing a non-target shRNA. Immunohistochemistry (IHC) analysis of residual tumors revealed a marked reduction of Ki-67-positive proliferating cells in doxycycline-treated tumors (Fig. 2B), consistent with the results of in vitro studies.

**Figure 2.**
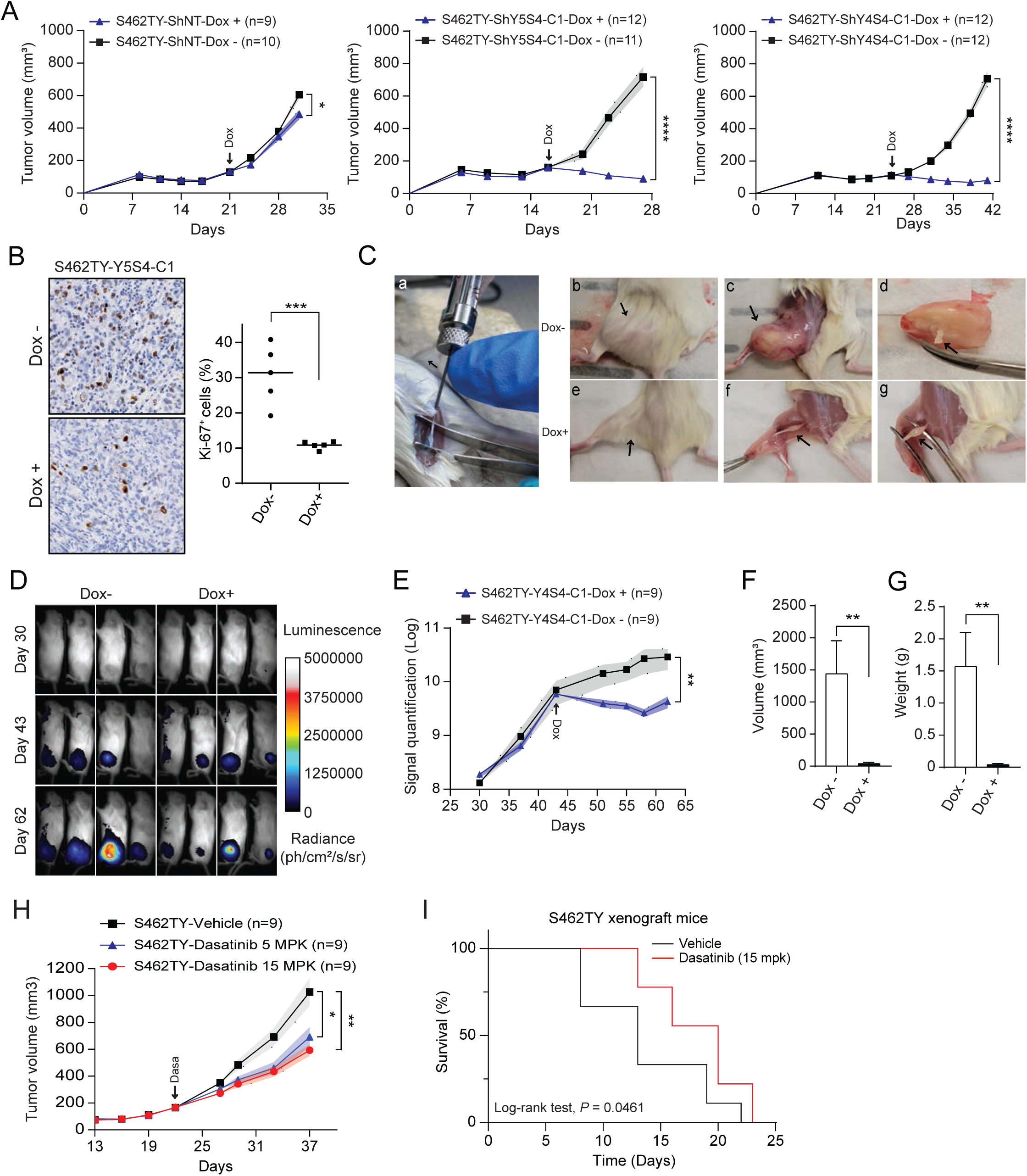
Targeting of YES/SRC inhibits MPNST tumor growth. (**A**-**B**) Ectopic MPNST model. S462TY control cells (shNT) or cell clones (Y5S4 and Y4S4) stably expressing *YES1*/*SRC* Dox-inducible shRNAs were grafted subcutaneously in NSG mice. Once tumors reached a volume of 120-150 mm^3^, the mice (n = 9-12 mice/group) were randomized and treated with vehicle or Dox (0.5 mg/mL in drinking water) for the indicated times. (**A**) Tumor growth was monitored bi-weekly. Data are means ± SEM (unpaired Student’s t-test). (**B**) IHC analysis of Ki-67 proliferation marker. Left, representative staining. Right, quantification of IHC staining (n=5, unpaired Student’s t-test). (**C**-**E**) Orthotopic MPNST model. S462TY cells (clone Y4S4) stably expressing Dox-inducible *YES1*/*SRC* shRNAs and luciferase were injected into the sciatic nerve of NSG mice. At day 44, the mice (n = 9 mice/group) were randomized and treated with vehicle or Dox for 3 weeks. Tumor growth was monitored by bioluminesce imaging (BLI). (**C**) Representative pictures: a) injection into the sciatic nerve; b and e) macroscopic appearance of tumors; c and f) images of intrasciatic tumors; d and g) dissected tumors shown with the sciatic nerve. (**D**) Representative bioluminescence images at day 30, 43, and 62 post-engraftment. (**E**) Quantification of tumor volume by BLI. Data are means ± SEM (unpaired Student’s t-test). (**F** and **G**) Final tumor volume (**F**) and weight (**G**) at the end of the study. Data are means ± SEM (unpaired Student’s t-test). (**H** and **I**) Efficacy studies of dasatinib. S462TY cells were implanted subcutaneously in NSG mice. Once tumors reached a volume of 120-150 mm^3^, the mice were randomized to treatment with vehicle or dasatinib (5 and 15 mg/kg) administered daily by oral gavage for the indicated times. (**H**) Tumor growth was monitored bi-weekly. Data are means ± SEM (n=9, unpaired Student’s t-test). (**I**) Kaplan-Meier analysis of overall survival (n=9 mice/group, log-ranked test). ns, not significant. \**P* < 0.05, ** *P* < 0.01, *** *P* < 0.001, **** *P* < 0.0001.

To evaluate the role of YES and SRC in a more clinically relevant setting, we established an orthotopic MPNST model by injecting luciferase-transfected S462TY cells expressing conditional *YES1* and *SRC* shRNAs into the sciatic nerve of NSG mice (Fig. 2C). Tumor growth was monitored longitudinally by bioluminescence imaging. Based on pilot studies defining the kinetics of tumor development, treatment with vehicle or doxycycline was initiated 44 days after injection. After 3 weeks of treatment, we observed a significant reduction in bioluminescent signal in the doxycycline group compared to vehicle controls (Fig. 2D and E). At sacrifice, tumors from doxycycline-treated mice exhibited markedly lower volumes and weights than control tumors (Fig. 2F and G), mirroring results from the subcutaneous model.

To explore the pharmacological potential of targeting YES/SRC in MPNST, we evaluated the antitumor efficacy of dasatinib in the S462TY xenograft model. NSG mice were implanted subcutaneously with S462TY cells, and treatment was initiated when tumors reached a volume of ∼150 mm^3^. The mice were randomized to receive vehicle or dasatinib at 5 or 15 mg/kg once daily for two weeks. Oral administration of dasatinib significantly slowed tumor growth with a 42% reduction in tumor volume observed at day 15 (Fig. 2H). Importantly, treatment with dasatinib significantly prolonged the overall survival of S462TY tumor-bearing mice (Fig. 2I). Together, our results indicate that dual targeting of YES/SRC activity can inhibit the growth of MPNST in preclinical models.

### YES/SRC signaling regulates oncogenic transcriptional programs and antigen processing and presentation pathway in MPNST

To identify downstream target genes of YES/SRC and explore the molecular consequences of their dual inhibition in MPNST, we analyzed the transcriptomes of control and YES/SRC-depleted S462TY cells by bulk RNA sequencing (RNA-seq). Principle component analysis (PCA) and hierarchical clustering revealed clear segregation between control and YES/SRC-depleted S462TY cells, with robust clustering of replicates within each group (Fig. 3A and B). Loss of YES and SRC induced broad transcriptional reprogramming of S462TY cells, with 2,358 genes significantly upregulated and 1,033 genes downregulated (fold change ≥ 2) relative to controls (Fig. 3C and Dataset EV1). Gene set enrichment analysis (GSEA) of differentially expressed genes (DEGs) revealed a marked downregulation of transcriptional programs associated with cell cycle progression, mitosis, DNA replication, E2F targets, and mitogenic signaling pathways (Fig. 3D-F and Tables EV1-EV3), consistent with a major role of YES/SRC signaling in sustaining MPNST cell proliferation. Notably, we observed a negative enrichment of RAS-ERK1/2 MAP kinase, PI3K/AKT, and mTORC1 signature genes, along with MYC target genes, underscoring the importance of YES/SRC in maintaining signaling pathways aberrantly activated in *NF1*-mutant MPNST (Yao et al., 2023). In addition, depletion of YES/SRC led to downregulation of canonical YAP/TAZ target genes, further supporting a functional link between the two signaling pathways (Fig. 3G). Thus, dual inactivation of YES and SRC disrupts multiple oncogenic signaling pathways central to MPNST pathogenesis.

**Figure 3.**
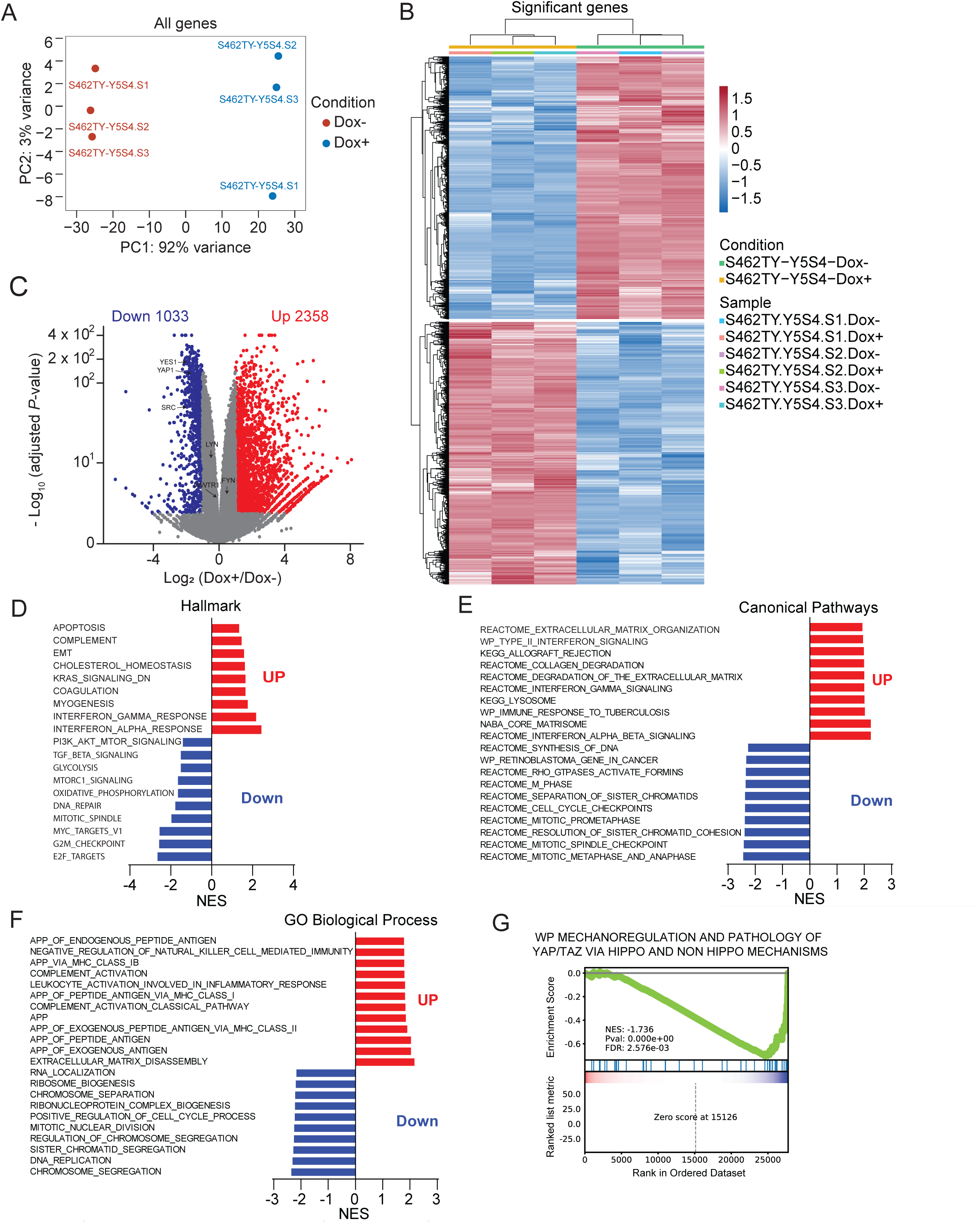
Targeting of YES/SRC reprograms oncogenic and immune signaling transcriptional networks in MPNST. S462TY cells (clone Y5S4) expressing Dox-inducible *YES1*/*SRC* shRNAs were treated with vehicle or 400 ng/ml Dox for 4 days. Whole transcriptomes were analyzed by RNA-seq. (**A**) PCA of all DEGs in control (red) and YES/SRC-depleted (blue) S462TY cells. Each dot corresponds to a single replicate. (**B**) Hierarchical clustering of DEGs. (**C**) Volcano plot of DEGs. Upregulated and downregulated genes with a log_2_(fold-change) ≥ 1 and *P* < 0.05 are represented as red and blue dots, respectively. (**D** - **G**) GSEA of DEGs. Enriched gene sets are expressed as normalized enrichment scores (NES). Red and blue bars correspond to gene sets upregulated or downregulated in YES/SRC-depleted S462TYcells, respectively. The top 10 altered gene sets are shown. (**D**) Hallmark gene sets. (**E**) Canonical pathways gene sets. (**F**) Gene Ontology Biological Process gene sets. (**G**) GSEA plot of Mechanoregulation and pathology of YAP/TAZ via Hippo and nonHippo mechanisms gene set.

Interestingly, depletion of YES/SRC also triggered a marked upregulation of gene signatures associated with interferon (IFN) signaling and antigen processing and presentation (APP) pathway (Fig. 3D-F and Tables EV1-EV3). Upregulated transcripts included MHC class I heavy chain genes and *B2M*, as well as genes involved in peptide trafficking and presentation, such as *TAPBP* and the endoplasmic reticulum chaperone *CALR* (Fig. 4A). In addition, markers of leukocyte activation, phagocytosis, and inflammatory signaling were elevated, indicative of broader remodeling of immune-related programs (Fig. EV6). To validate these transcriptomic findings, we measured MHC class I surface expression by flow cytometry after genetic or pharmacological inhibition of YES/SRC activity in S462TY cells. Doxycycline-induced shRNA-mediated depletion of YES/SRC significantly increased HLA class I expression in both clones of S462TY cells (Fig. 4B). Similarly, treatment of S462TY cells with 100 nM dasatinib for 72 h upregulated HLA class I levels to an extent comparable to interferon gamma (IFN-γ) stimulation (Fig. 4C). The effect was reproduced with structurally distinct SFK inhibitors, further supporting the specific involvement of YES and SRC kinases (Fig. 4D). To begin dissecting the underlying mechanisms, we examined the impact of YES/SRC inhibition on upstream signaling events known to regulate MHC class I gene expression (Kobayashi & van den Elsen, 2012). Both genetic depletion and pharmacological inhibition of YES/SRC led to a robust increase in STAT1 Tyr701 phosphorylation associated with elevated expression of the MHC class I transactivator NLRC5 (Fig. 4E). Collectively, these results suggest that YES/SRC signaling not only drives MPNST proliferation but may also repress antitumor immune responses.

**Figure 4.**
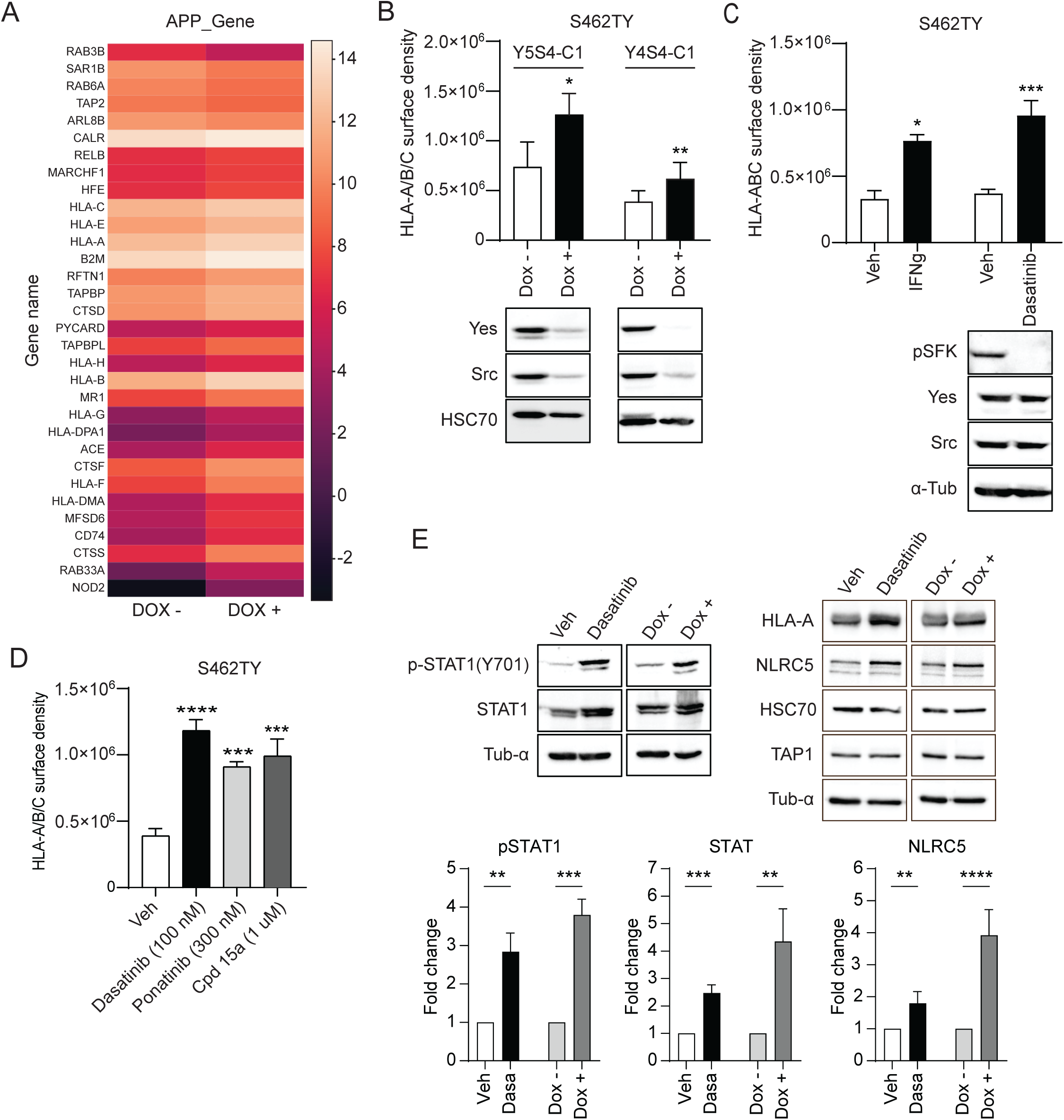
Inhibition of YES/SRC signaling upregulates APP pathway genes and MHC class I expression in MPNST cells. (**A**) Heat map showing differential expression of APP-related genes upon Dox-induced depletion of YES/SRC in S462TY cells. Genes were selected from the Gene Ontology Biological Process Antigen processing and presentation gene set. The color corresponds to the fold-change in expression. (**B**) S462TY clones expressing Dox-inducible *YES1*/*SRC* shRNAs were treated with vehicle or 400 ng/ml Dox for 4 days. HLA-A/B/C surface expression was analyzed by flow cytometry. Data are means ± SEM (n=3, unpaired Student’s t-test). Bottom, representative western blots. (**C**) S462TY cells were treated with vehicle or either 40 ng/ml IFN-γ or 100 nM dasatinib for 72 h. Quantification of HLA-A/B/C surface expression. Data are means ± SEM (n=3, unpaired Student’s t-test). Bottom, representative western blots. (**D**) S462TY cells were treated with vehicle or either 100 nM dasatinib 300 nM ponatinib or 1 μM Compound 15a for 72 h. HLA-A/B/C surface expression was analyzed by flow cytometry. Data are means ± SEM (n=3, one-way ANOVA with Dunnett’s post hoc test). (**E**) Analysis of MHC class I expression signaling pathway. S462TY cells were treated with dasatinib for 72 h or depleted of YES/SRC for 4 days. Cell lysates were analyzed by western blotting. Top, representative western blots. Bottom, quantification of immunoblotting data normalized to a-tubulin. Data are means ± SEM (n=3, unpaired Student’s t-test).

### Phosphotyrosine proteomics reveals a YES/SRC-dependent network regulating MAP kinase and immune signaling in MPNST

To further delineate the YES/SRC signaling network in MPNST and identify candidate substrates and downstream effectors, we profiled the global phosphotyrosine proteome of S462TY cells under two complementary perturbations: (1) long-term (96 h) genetic depletion of YES/SRC proteins by shRNA; and (2) acute (1 h) pharmacological inhibition of SFK activity with dasatinib (Fig. 5A). Phosphotyrosine-containing peptides were enriched by anti-phosphotyrosine immunoprecipitation and analyzed by LC-MS/MS on a Ascend Orbitrap instrument. Label-free quantification, normalized to changes in protein abundance, was used to define differentially regulated sites (absolute log2(fold change) ≥ 1, *P* ≤ 0.05). Upon depletion of YES/SRC, we quantified 187 phosphotyrosine sites on 112 proteins, of which 29 were significantly downregulated and 21 upregulated (Fig. 5A and Dataset EV2). Dasatinib treatment yielded 125 sites on 84 proteins, with 24 sites decreased and 2 increased (Fig. 5B and Dataset EV3). Activating tyrosines of FYN/LYN/YES1/SRC (peptide sequence identical) were strongly dephosphorylated, consistent with direct inhibition of SFK catalytic activity by dasatinib. Across both perturbations, the phosphoproteome converged on two major signaling axes: the canonical SFK-adaptor protein-MAPK cascade and a second tier of cytokine/JAK-STAT components linked to MHC class I antigen processing. Prominent shared changes included dephosphorylation of multiple sites on CRKL, decreased ERK2/MAPK1 Y187 and JNK2/MAPK9 Y185, and loss of phosphorylation of TYK2 Y292. GO term enrichment analysis (Fig. 5C) highlighted MAPK cascade, Intracellular signal transduction and JNK cascade pathways, consistent with coordinated downregulation of SFK-adaptor protein-MAPK signaling nodes. In parallel, IFN-JAK-STAT signaling terms emerged, driven by changes in TYK2, JAK2, STAT3 and IFITM3 tyrosine phosphorylation. The overlap between the two datasets therefore defines a shared core response to inhibition of YES/SRC signaling.

**Figure 5.**
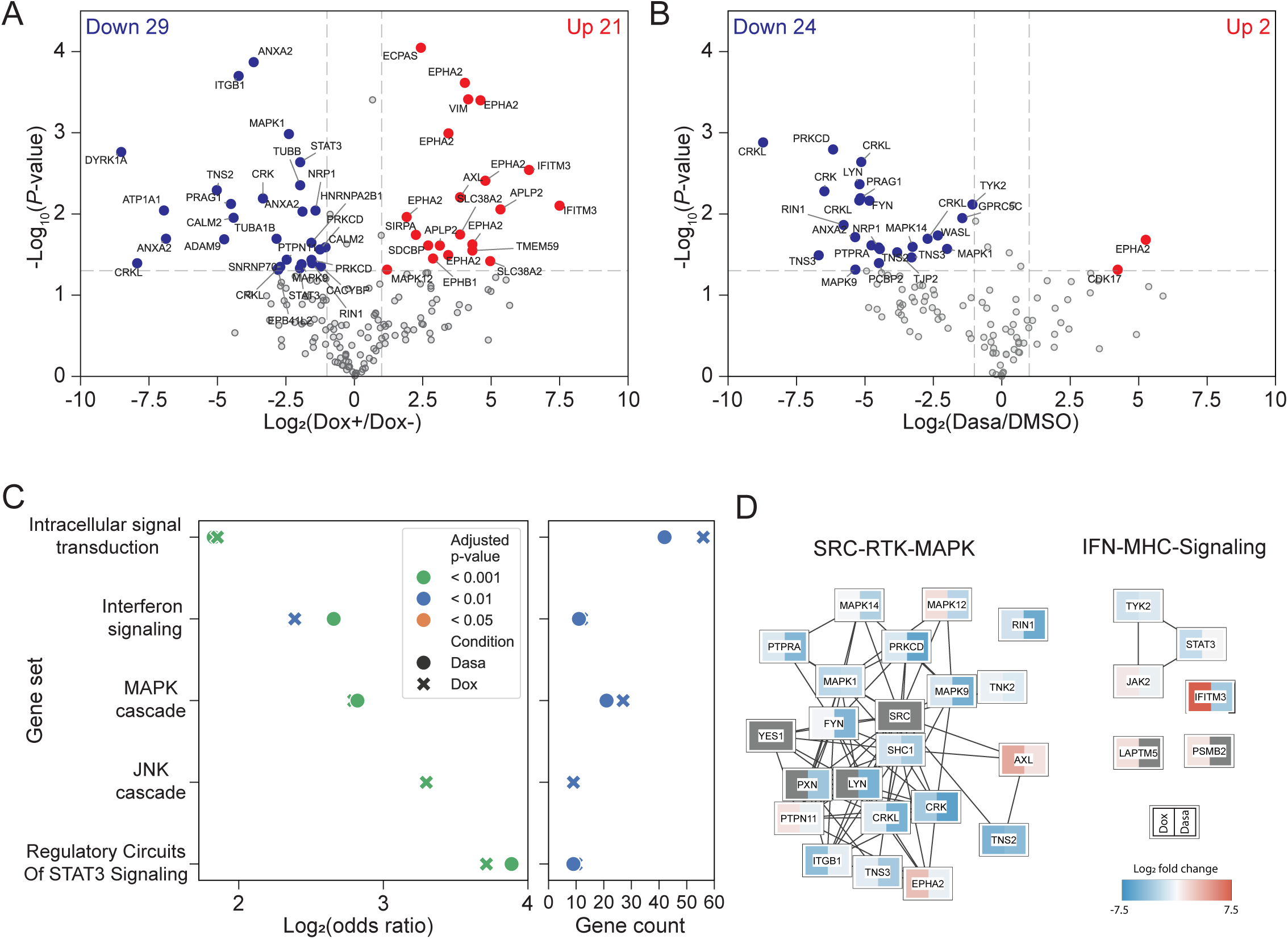
Global analysis of the YES/SRC-dependent tyrosine phosphoproteome in MPNST. (**A** and **B**) Volcano plot of phosphotyrosine sites quantified after Dox-induced shRNA-mediated depletion of YES/SRC (clone Y5S4) (**A**) or dasatinib treatment (**B**). Each point represents a phosphotyrosine site. Dashed lines indicate the fold-change and significance cutoffs. Significantly downregulated and upregulated sites are highlighted in blue and red, respectively. (**C**) Over-representation analysis of proteins containing significantly regulated phosphotyrosine sites. Enrichment is shown as log_2_(odds ratio) (left) and gene count per term (right). Symbols denote perturbation and colors indicate adjusted *P*-value bins. (**D**) Interaction networks of phosphoregulated proteins connecting YES/SRC signaling to receptor tyrosine kinase (RTK)–MAPK (left) and IFN–MHC (right) signaling modules. The networks were generated using STRING. Nodes represent proteins, while edges denote curated protein–protein interactions. Node tiles report log₂ fold-change for Dox (left half) and dasatinib (right half) using the indicated color scale; gray denotes not quantified/unchanged in the corresponding condition.

The canonical adaptors CRK and CRKL were among the most affected substrates: multiple sites within the CRKL C-terminal region and CRK Y221 lost phosphorylation by up to 9-fold. These adaptor proteins play a major role in the regulation of MAPK pathways by integrating signals from different sources and often synergizing with activated RAS to drive cancer progression (Birge et al, 2009). Consequently, we observed a decrease in phosphorylation of the TXY activation motifs of ERK2 (MAPK1 Y187), JNK2 (MAPK9 Y185) and p38α (MAPK14 Y182) after inhibition of YES/SRC signaling. In parallel, the SRC-proximal non-receptor kinase TNK2 (Y827) and focal adhesion-associated scaffolds TNS2 (Y483) and TNS3 (Y354 and Y354) were markedly dephosphorylated. Both datasets also revealed modulation of IFN-JAK-STAT and APP components. Notably, YES/SRC depletion led to significant dephosphorylation of STAT3 Y705, the canonical activating site, together with strong upregulation of IFITM3 Y20 and increased phosphorylation of SIRPA Y496.

To place these events in their signaling context, we constructed an interaction network of phosphoregulated proteins emphasizing protein kinases, adaptors, and scaffolds bridging the SFK/receptor tyrosine kinases/MAPK and IFN-JAK-STAT signaling axes (Fig. 5D). This network revealed a dense module of proteins propagating adhesion- and growth factor-derived signals into MAPKs (SRC, YES1, FYN, LYN, EPHA2, AXL, TNK2, CRK, CRKL, MAPK1, MAPK9, MAPK12, MAPK14, PRKCD, PTPN11, PTPRA, RIN1, TNS2, TNS3, PXN, ITGB1) and components linked to IFN-induced antigen processing and MHC class I expression (JAK2, TYK2, STAT3, IFITM3, PSMB2, LAPTM5). Collectively, these data show that YES/SRC coordinately signal to oncogenic MAPK and immune-modulatory IFN-JAK-STAT phospho-networks, providing a mechanistic framework for how SFK inhibition can simultaneously restrain tumor cell proliferation and reshape the antigen-presentation landscape in MPNST cells.

### *YES1* is overexpressed in human MPNST

Given the limited clinical annotation available for existing MPNST patient cohorts, we interrogated a large sarcoma transcriptomic dataset from the Centre Léon Bérard to compare *YES1* and *SRC* expression between MPNST and benign soft tissue and bone tumors. We found that *YES1*, but not *SRC*, is expressed at a significantly higher level in MPNST relative to benign tumors (Fig. 6A and B). Consistent with a pathogenic role, high *YES1* expression predicts shorter overall survival in the TCGA cohort of sarcoma patients (Lapouge & Meloche, 2023).

**Figure 6.**
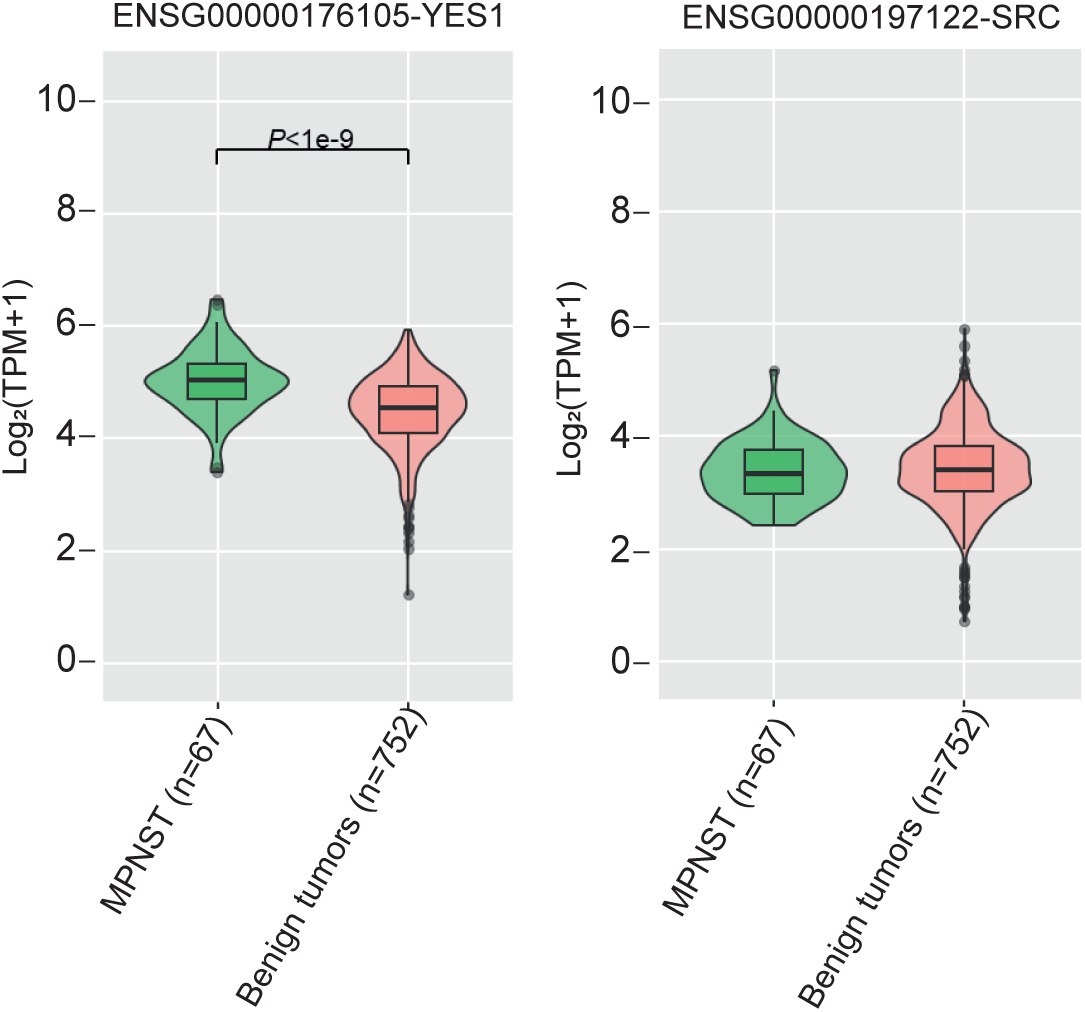
Overexpression of *YES1* in human MPNST. Comparative expression of *YES1* and *SRC* genes in MPNST and benign soft tissue and bone tumors of distinct histological subtypes. Violin and boxplot RNA-seq expression values are represented as log2 of transcript per million (TPM) +1. The number of samples is indicated in parentheses. Statistical significance was assessed by unpaired Student’s t-test.

## DISCUSSION

Despite advances in defining the genomic landscape and molecular pathogenesis of MPNST, this malignancy remains one of the most lethal soft-tissue sarcomas with very limited therapeutic modalities (Martin et al., 2020; Somaiah et al, 2024; Yao et al., 2023). While surgery remains the mainstay of treatment for localized MPNST, there is no effective systemic therapy for advanced or metastatic disease. In this context, our study identifies the SFKs YES and SRC as redundant, non-oncogene addictions in MPNST and uncovers a dual oncogenic and immune-regulatory vulnerability with therapeutic potential.

Prior studies have implicated Hippo-YAP/TAZ pathway dysregulation in Schwann cell transformation and MPNST (Isfort et al., 2019; Velez-Reyes et al., 2021; Wu et al., 2018). We first confirmed that YAP/TAZ signaling is broadly required for the proliferation of benign plexiform neurofibroma and malignant MPNST cell lines, and that YAP/TAZ protein levels are increased in human MPNST compared with benign nerve sheath tumors. Our findings extend previous observations and position YAP/TAZ as central effectors across the neurofibromatosis spectrum. Given the established role of the SFKs YES and SRC as upstream regulators of Hippo-YAP/TAZ signaling, we next assessed their functional importance in MPNST biology. Using complementary genetic and pharmacological approaches, we found that dual inhibition of YES and SRC signaling markedly impaired the proliferation of multiple NF1-mutant MPNST cell lines. Importantly, CRISPR/Cas9-mediated gene editing demonstrated that concurrent loss of *YES1* and *SRC* genes is not tolerated, indicating that these kinases are redundantly required for MPNST viability. These findings reinforce the rationale for concomitant pharmacological targeting of both tyrosine kinases in MPNST. The in vivo relevance of this dependency was demonstrated in complementary mouse xenograft models of MPNST. Conditional depletion of YES/SRC in S462TY tumor cells resulted in near-complete arrest of tumor growth in both subcutaneous and orthotopic sciatic nerve models, associated with a marked reduction of cell proliferative. Treatment of established S462TY tumors with dasatinib significantly delayed tumor growth and improved overall survival.

Integrated transcriptomic and phosphotyrosine proteomic analyses revealed how YES/SRC signaling regulates core oncogenic networks and sustains cell proliferation in MPNST. Dual depletion of YES/SRC elicited broad transcriptional reprogramming, with coordinated repression of cell cycle, DNA replication, mitosis, and E2F programs, and downregulation of oncogenic RAS–ERK1/2 MAPK, PI3K/AKT, mTORC1, MYC, and YAP/TAZ targets. These signaling pathways correspond closely to those aberrantly activated in NF1-mutant and PRC2-deficient MPNST (Kim & Pratilas, 2018; Somatilaka et al, 2022; Suppiah et al., 2023; Yao et al., 2023). Phosphotyrosine profiling converged on a core network characterized by dephosphorylation of the adaptors CRK/CRKL, inactivation of ERK1/2, JNK, and p38 MAPKs, and altered phosphorylation of RTK-associated scaffolds. Notably, despite compensatory hyperphosphorylation of several RTKs, these receptors are uncoupled from MAPK modules when YES/SRC signaling is inhibited, suggesting that YES/SRC provide an essential scaffolding function that routes diverse upstream inputs into mitogenic pathways. Targeting this convergence node may therefore help overcome pathway redundancy and adaptive resistance that limit the efficacy of single-agent therapies in MPNST and other sarcomas.

A major and unexpected finding is that YES/SRC signaling also regulates the APP pathway in MPNST cells. Transcriptomic analysis revealed strong enrichment of IFN and APP gene sets upon YES/SRC depletion, including upregulation of MHC class I, peptide-loading components, and chaperones. Consistent with these changes, genetic or pharmacologic YES/SRC inhibition increased HLA class I surface expression on MPNST cells. Mechanistically, we observed induction of STAT1 Tyr701 phosphorylation and upregulation of NLRC5, the master transactivator of MHC class I genes, concomitant with decreased STAT3 Tyr705 phosphorylation and modulation of IFITM3 Tyr20 and TYK2 phosphorylation. These data support a model in which YES/SRC activity dampens an IFN–STAT1–NLRC5 axis and maintains low MHC class I expression, thereby facilitating immune evasion (Cui et al, 2025; Kobayashi & van den Elsen, 2012; Rebe & Ghiringhelli, 2019). These observations have important therapeutic implications. Clinical and translational studies consistently characterize MPNST as immune-cold tumors with low T-cell infiltration and limited responses to immune checkpoint inhibitors (Lee et al., 2006; Lingo et al, 2025; Paudel et al., 2023). Our data suggest that YES/SRC inhibition may convert a fraction of MPNSTs from immune-cold to a more inflamed, MHC class I–high state, potentially enhancing antigen presentation to T cells. In this regard, YES/SRC inhibitors could be leveraged as a priming strategy to increase the efficacy of immune checkpoint blockade in NF1-associated and sporadic MPNST.

In summary, we identify the tyrosine kinases YES/SRC as actionable, candidate therapeutic targets in MPNST. Dual inhibition of YES/SRC suppresses multiple mitogenic signaling networks while simultaneously relieving repression of immune-regulatory pathways, thereby addressing two major barriers to effective therapy in this disease. The observation that *YES1* is significantly overexpressed in MPNST relative to benign soft-tissue tumors, and that high *YES1* expression predicts shorter overall survival in the TCGA sarcoma cohort further support the translational relevance of our findings. Single-agent dasatinib did not show significant clinical activity in unselected patients with high-grade advanced sarcomas (Schuetze et al, 2016). Future studies with next-generation, more selective YES/SRC inhibitors, as well as rational combination with targeted agents or immunotherapies will be essential to define the clinical potential of targeting YES/SRC in MPNST.

## METHODS

### Reagents and antibodies

Dasatinib (D-3307) was purchased from LC Laboratories. Ponatinib (S1490) was from Selleck Chemicals. Compound 15a was synthesized in-house as described (Du et al, 2020). Doxycycline (Dox) was purchased from Sigma-Aldrich. Matrigel basement membrane matrix was from Corning.

Commercial antibodies were obtained from the following suppliers: anti-YES (610375) from BD Biosciences-Pharmingen; anti-SRC (32G6), anti-phospho-SFK(Y416) (2101), anti-YAP (1A12), anti-YAP (D8H1X for immunohistochemistry (IHC), anti-YAP/TAZ (D24E4 for IHC detection of both YAP and TAZ), anti-STAT1 (9172), anti-phospho-STAT1(Y701) (7649), anti-NLRC5 (E1E9Y), and anti-TAP1 (E4T4F) from Cell Signaling Technology; anti-HLA-A (ab52922) and anti-α-tubulin (ab18251) from abcam; anti-HSC70 (sc-7298) from Santa Cruz Biotechnology; anti-Ki-67 (CRM325A) from Biocare Medical.

### Cell Culture

Human plexiform neurofibroma cell lines ipNF05.5 and ipNF95.6, and human MPNST cell lines sNF96.2 and sNF02.2 were obtained from American Type Culture Collection. The human S462TY cell line was kindly provided by Dr Timothy Cripe (Ohio State University). All cell lines were cultured in Dulbecco’s modified Eagle’s medium (DMEM) supplemented with 10% fetal bovine serum (FBS), glutamine, and 1% penicillin-streptomycin in a humidified atmosphere of 5% CO_2_ and 95% air at 37^0^C. The cells were regularly tested for mycoplasma contamination.

### RNA interference and CRISPR/Cas9 gene editing

For transient RNA interference experiments, cells were transfected with SMARTpool siRNAs (Dharmacon) targeting *YES1* (L-003184-00-0005), *SRC* (L-003175-00-0050), *YAP1* (L-012200-00-0050) or *WWTR1* (L-016083-00-0050) using Lipofectamine RNAiMAX (Thermo Fisher Scientific) according to the manufacturer’s instructions.

For inducible shRNA experiments, the plasmids encoding Dox-inducible *YES1* and *SRC* shRNAs were generated by subcloning *YES1*- and *SRC*-specific shRNA sequences from the TRC1 shRNA library (Sigma-Aldrich) into the pLKO-Tet-On lentiviral expression vector. The following shRNA sequences were used: TRCN0000010006 (*YES1* S4), TRCN0000121230 (*YES1* S5),

TRCN0000038150 (*SRC* S4), and TRCN0000038149 (*SRC* S5). Lentiviral infections of S462TY cells were performed as previously described (Guegan et al., 2022). Stable populations of infected cells were selected with 1 µg/mL puromycin for 4 days. Clonal cell lines were isolated by limiting dilution. Conditional depletion of SRC and YES was induced by addition of 400 ng/ml Dox to the culture medium.

For CRISPR/Cas9 gene editing, the plasmids pLentiCRISPRv2-YES1 (sgRNA sequence 5’-AGGTGGTGTCACTATATTTG-3’) and pLentiCRISPRv2-SRC (sgRNA sequence 5’-TCCGTGACTCATAGTCATAG-3’) were purchased from GenScript and validated internally. After lentiviral infection, stable populations of S462TY cells were selected with 1 µg/mL puromycin. Single-cell derived clones were isolated by limiting dilution.

### Western blot analysis

Cell lysis and western blot analysis were performed as described previously (Voisin et al, 2024). Immunoreactive bands were detected by enhanced chemiluminescence (ECL Plus, GE Healthcare) and visualized with a ChemiDoc imaging system (Bio-Rad). For dot blot analysis, single-cell S462TY clones were seeded in 96-well plates and cultured until reaching ∼ 80% confluence. Then, 10 μL of lysis buffer was added directly to each well, and 4 μL of the lysate was spotted onto a dry nitrocellulose membrane and allowed to air-dry. Membranes were blocked in blocking buffer (Tris-buffered saline containing 5% non-fat dry milk and 0.1% Tween 20) for 1 h at room temperature, followed by incubation with primary antibodies overnight at 4 °C. Immunoreactive spots were visualized by enhanced chemiluminescence.

### Cell proliferation assay

Cell proliferation was assessed using the WST-1 colorimetric assay (Roche) according to the manufacturer’s instructions. Dose-response curves were analyzed by nonlinear regression using GraphPad Prism.

### Gene expression profiling and pathway enrichment analysis

The transcriptome of S462TY MPNST cells conditionally depleted of YES/SRC expression was analyzed by RNA sequencing. S462TY cells expressing Dox-inducible *YES1*/*SRC* shRNAs were treated with vehicle (n=3) or 400 ng/ml Dox (n=3) for 4 days. Total RNA was extracted using RNAeasy purification kit (Qiagen) and RNA integrity was assessed with the Agilent 2100 Bioanalyzer. Strand-specific RNA libraries were prepared as previously described (Voisin et al., 2024) and sequenced on an Illumina NextSeq500 sequencer, generating around 15 million single-end reads per sample. Adapter sequences and low-quality bases in the resulting FASTQ files were removed using Trimmomatic version 0.35 (Bolger et al, 2014). Reads were aligned to the human reference genome GRCh38 (from Gencode version 37 based on Ensembl 103) using STAR version 2.7.1a (Dobin et al, 2013). DESeq2 (Love et al, 2014) was used to normalize gene readcounts and compute differential expression between experimental conditions. Differentially expressed genes (DEGs) with an adjusted P value < 0.05 were considered significant. Pathway enrichment analysis of pre-ranked gene lists was performed using the GSEApy software v1.0.4 (Fang et al, 2023) and the human Molecular Signature Database of annotated gene sets (Liberzon et al, 2015). Gene sets were filtered using an absolute value of the normalized enrichment score (NES) > 1.4 and false discovery rate (FDR) < 0.1.

### Flow cytometry analysis of HLA class I expression

Surface expression of HLA class I (HLA-A/B/C) was quantified using the Qifikit (Dako) according to the manufacturer’s instructions. The cells were stained on ice in phosphate-buffered saline (PBS) supplemented with 2% FBS to minimize receptor internalization and nonspecific binding. Staining was performed using the primary anti-HLA-A/B/C monoclonal antibody W6/32 (ThermoFisher), followed by Alexa Fluor 488–conjugated anti-mouse IgG secondary antibody. For each condition, 10,000 single, live cells were acquired on a Yeti flow cytometer (Propel Labs). Gating strategy included exclusion of debris and doublets based on forward and side scatter (FSC/SSC) parameters, followed by singlet discrimination to ensure accurate analysis of individual viable cells. Mouse IgG2a kappa isotype control (clone eBM2a, ThermoFisher) was included in each experiment to account for nonspecific antibody binding. Unstained samples were analyzed during initial assay optimization to validate that cellular autofluorescence did not interfere with specific signal detection under the selected acquisition settings. Quantification of HLA class I surface expression was performed by converting the median fluorescence intensity (MFI) into the number of antibody-binding sites per cell using the calibration beads provided in the Qifikit. Data were analyzed using the FlowJo software (version 10.10).

### Quantitative tyrosine phosphoproteomic analysis

Analysis of the YES/SRC-dependent tyrosine phosphoproteome in MPNST cells was performed using complementary genetic and pharmacological perturbation conditions. Biological triplicates of S462TY cells expressing Dox-inducible *YES1*/*SRC* shRNAs were treated with vehicle or 400 ng/ml Dox for 4 days. In parallel, parental S462TY cells (n=3) were treated with vehicle or 100 nM dasatinib for 1 h. The cells were lysed in Tris-HCl (pH 8.1), 8 M urea, 5 mM TCEP, 20 mM chloroacetamide buffer and protein extracts were digested with sequencing-grade trypsin (1 μg). Phosphotyrosine-containing peptides were enriched using pY100 anti-phosphotyrosine antibody-conjugated agarose beads (Cell Signaling Technology) according to the manufacturer’s protocol. Enriched peptides were analyzed by nano liquid chromatography-tandem mass spectrometry (LC-MS/MS) using a Thermo Scientific Vanquish Neo coupled to an Thermo Ascend Orbitrap. Samples were loaded on a Thermo Pepmap C18 precolumn and eluted from an IonOpticks Aurora 25-cm column with a 120 min gradient. MS survey scans were acquired at 120K resolution and a 50ms injection time. Tandem mass spectra were acquired at 30K resolution and 59 ms injection time. Proteome samples were analyzed by nanoLC-MS/MS using a Thermo Scientific EasynLC1200 coupled to an Thermo Exploris Orbitrap with the FAIMS interface. Samples were loaded on a Optimize Technologies C4 precolumn and eluted from a homemade 25-cm column packed with Phenomenex Jupiter C18 with a 120 min gradient as described (Wu et al, 2022). MS survey scans were acquired at 120K resolution with a 50 ms injection time. Tandem mass spectra were acquired at 30K resolution and 59 ms injection time. Data were processed with PEAKS 10.6 and searched against the UniProt human reference proteome (20,359 entries). Carbamidomethylation of cysteines was selected as fixed modifications. M oxidation, NQ deamidation and STY phosphorylation were selected as variable modifications. Peptide and protein identifications were filtered using a 1% FDR. PEAKS label-free quantification workflow was used to extract peptide features, and differential analysis of phosphorylation was done using MSstatsPTM v2.4.1 (Kohler et al, 2023). Because of the long treatment time in the *YES1*/*SRC* shRNAs condition, the changes in phosphorylation ratios were normalized to protein change ratios. GO term enrichment analysis was performed using the gProfiler web service (Raudvere et al, 2019). Protein interaction networks were extracted from STRING v12 (Szklarczyk et al, 2025) and visualized with Cytoscape v3.10.4 (Shannon et al, 2003).

### Animal procedures

NSG mice were obtained from The Jackson Laboratory. The mice were housed in specific pathogen–free (SPF) conditions in filter-topped isolator cages, with ad libitum access to food and water. All animal procedures were approved by the institutional animal care committee and conducted in accordance with the guidelines of the Canadian Council on Animal Care and applicable national regulations. For subcutaneous transplantation experiments, 6–8-week-old female NSG mice were injected subcutaneously in the right flank with 5 × 10⁶ cells resuspended in PBS with 50% Matrigel. Once tumors reached a volume of 120–150 mm³, the mice were randomized to the different treatment groups. Mice injected with S462TY cells harboring Dox-inducible *YES1*/*SRC* shRNAs were treated with vehicle or doxycycline (0.5 mg/mL) in drinking water. For pharmacological studies, mice injected with parental S462TY cells were randomized to treatment with vehicle or dasatinib (15 mg/kg) administered daily by oral gavage for the indicated times. Tumor growth was measured bi-weekly with a Vernier caliper and tumor volumes were calculated using the formula: V = (length × width^2^)/2. For analysis of overall survival, the mice were monitored until they reach humane endpoints. Survival was estimated using the Kaplan-Meier method based on the proportion of animal alive at each time point.

For orthotopic transplantation experiments, 3 × 10⁵ luciferase-labeled S462TY cells harboring Dox-inducible *YES1*/*SRC* shRNAs (clone Y4S4) were injected directly into the sciatic nerve of female NSG mice. The mice were anesthetized and placed in lateral recumbency. A small incision was made through the skin and fascia lata of the thigh to expose the sciatic nerve, taking care not to damage the underlying muscle. Cells were injected in a volume of 3 µL using a Hamilton syringe fitted with a 34-gauge needle. The incision was closed with absorbable sutures, and mice received post-operative analgesia and routine monitoring in accordance with institutional animal care guidelines. Tumor growth was monitored by bioluminescence imaging (BLI). Imaging was performed twice a week throughout the study with the injection of 150 mg/kg of fresh sterile D-Luciferin (MediLumine). Images were obtained 15 min after the intraperitoneal injections of D-Luciferin using the LabeoTech OiS300 In Vivo Imaging System (Labeo Technologies). Signal normalization and analysis was done automatically for all time points using ImageJ (version 2.1.0/1.53d) macros and expressed in radiance (photons·s^−1^·sr^−1^·cm^−2^) integrated density (area x mean intensity).

### Histology and immunohistochemistry analysis

Tumors were fixed for at least 24 h in 10% formalin, embedded in paraffin, and sliced in 4-mm thin sections. Tumor sections were mounted on glass slides and stained with H&E using established protocols. For immunohistochemistry (IHC) analysis, FFPE tissue sections were processed as described previously (Voisin et al., 2024) and stained with anti-Ki-67 antibody.

### MPNST tissue microarray construction and analysis

All patient provided informed consented for use of their tissue for research as part of the Interdisciplinary Health Research Team in Musculoskeletal Neoplasia biobank. The project was reviewed and approved by the MUHC institutional research ethics board. To construct the tissue microarrays (TMAs), we searched the biobank for all diagnosed cases, both biopsies and resections, of MPNST, atypical neurofibromatous neoplasm of uncertain biologic potential (ANNUBP), and neurofibromas (NF). Tumor tissue from 26 patients could be retrieved, including MPNST from 19 patients, ANNUBP from 12 patients, and NF from 11 patients. Of these 26 patients, 16 had NF1-associated tumors and 10 had sporadic tumors. Three patients had multiple or recurrent MPNSTs. A 2-mm core was punched out of the paraffin blocks to construct a 12 x 7 microarray. Tonsil and normal peripheral nerve tissues were incorporated as controls. TMAs were stained with the following primary antibodies: YAP (D8H1X; 1/100) and YAP/TAZ (D24E4; 1/50). Antibody conditions were optimized and validated on full tumor sections prior to TMA analysis. TMA slides were independently scored by two pathologists using a semi-quantitative method. Total (nuclear + cytoplasmic) staining intensity was scored and reported as the percentage of cells with significant staining. Membrane staining for phospho-SFK on TMAs was non-interpretable and therefore excluded from further analysis.

### Analysis of tumor clinical specimens

Expression of *YES1* and *SRC* transcripts in MPNST and benign tumors was analyzed using the large transcriptomic dataset from the Centre Léon Bérard in Lyon (Macagno et al, 2022).

### Statistical analyses

All statistical analyses were performed using GraphPad Prism (version 9.5.1) software. A *P* value < 0.05 was considered statistically significant. Data distribution was assumed to be normal, although normality was not formally tested. Statistical test used for each data set is indicated in the figure legend.

## DATA AVAILABILITY

RNA-seq data of S462TY cells have been deposited in the Gene Expression Omnibus (GEO) under the accession number GSE315714. The MS phosphoproteomics data have been deposited to the ProteomeXchange Consortium via the PRIDE partner repository with the dataset identifier PXD073130. All other data associated with this study are included in the main text or supplementary material.

## ACKNOWLEDGMENTS

We thank T. Cripe for the S462TY cell line. We thank Éric Bonneil for proteomic analysis, Raphaelle Lambert for RNA-seq, Patrick Gendron for advice on bioinformatic analyses, and Julie Hinsinger for histology assistance. E. Zereg is recipient of a studentship from the Fonds de recherche du Québec-Santé (FRQS). This study was supported by an Impact Grant (705039) from the Canadian Cancer Society Research Institute and by an Ad Hoc Project Grant (2023-02) from IRICoR through the Centres of Excellence for Commercialization and Research to S.M. The Institute for Research in Immunology and Cancer (IRIC) received infrastructure support from IRICoR, the Canadian Foundation for Innovation, FRQS, and the GTP program funded in part by Genome Canada and Génome Québec.

## Notes

### Competing Interest Statement

The authors have declared no competing interest.

## REFERENCES

Birge RB, Kalodimos C, Inagaki F, Tanaka S (2009) Crk and CrkL adaptor proteins: networks for physiological and pathological signaling. Cell Commun Signal 7: 13

Bolger AM, Lohse M, Usadel B (2014) Trimmomatic: a flexible trimmer for Illumina sequence data. Bioinformatics 30: 2114–2120

Cai Z, Tang X, Liang H, Yang R, Yan T, Guo W (2020) Prognosis and risk factors for malignant peripheral nerve sheath tumor: a systematic review and meta-analysis. World J Surg Oncol 18: 257

Cui Y, Qiu T, Wang J, Liu X, Luo L, Yin J, Zhi X, Wang W, Feng G, Wu C et al (2025) IFITM3 enhances immunosensitivity via MHC-I regulation and is associated with the efficacy of anti-PD-1/-L1 therapy in SCLC. Mol Cancer 24: 187

Dobin A, Davis CA, Schlesinger F, Drenkow J, Zaleski C, Jha S, Batut P, Chaisson M, Gingeras TR (2013) STAR: ultrafast universal RNA-seq aligner. Bioinformatics 29: 15–21

Du G, Rao S, Gurbani D, Henning NJ, Jiang J, Che J, Yang A, Ficarro SB, Marto JA, Aguirre AJ et al (2020) Structure-Based Design of a Potent and Selective Covalent Inhibitor for SRC Kinase That Targets a P-Loop Cysteine. J Med Chem 63: 1624–1641

Fang Z, Liu X, Peltz G (2023) GSEApy: a comprehensive package for performing gene set enrichment analysis in Python. Bioinformatics 39: btac757

Guegan JP, Lapouge M, Voisin L, Saba-El-Leil MK, Tanguay PL, Levesque K, Bregeon J, Mes-Masson AM, Lamarre D, Haibe-Kains B et al (2022) Signaling by the tyrosine kinase Yes promotes liver cancer development. Sci Signal 15: eabj4743

Isfort I, Elges S, Cyra M, Berthold R, Renner M, Mechtersheimer G, Aman P, Larsson O, Ratner N, Hafner S et al (2019) Prevalence of the Hippo Effectors YAP1/TAZ in Tumors of Soft Tissue and Bone. Sci Rep 9: 19704

Kantarjian H, Jabbour E, Grimley J, Kirkpatrick P (2006) Dasatinib. Nat Rev Drug Discov 5: 717–718

Kim A, Pratilas CA (2018) The promise of signal transduction in genetically driven sarcomas of the nerve. Exp Neurol 299: 317–325

Kobayashi KS, van den Elsen PJ (2012) NLRC5: a key regulator of MHC class I-dependent immune responses. Nat Rev Immunol 12: 813–820

Kohler D, Tsai TH, Verschueren E, Huang T, Hinkle T, Phu L, Choi M, Vitek O (2023) MSstatsPTM: Statistical Relative Quantification of Posttranslational Modifications in Bottom-Up Mass Spectrometry-Based Proteomics. Mol Cell Proteomics 22: 100477

Lamar JM, Xiao Y, Norton E, Jiang ZG, Gerhard GM, Kooner S, Warren JSA, Hynes RO (2019) SRC tyrosine kinase activates the YAP/TAZ axis and thereby drives tumor growth and metastasis. J Biol Chem 294: 2302–2317

Lapouge M, Meloche S (2023) A renaissance for YES in cancer. Oncogene 42: 3385–3393

Lee PR, Cohen JE, Fields RD (2006) Immune system evasion by peripheral nerve sheath tumor. Neurosci Lett 397: 126–129

Lee W, Teckie S, Wiesner T, Ran L, Prieto Granada CN, Lin M, Zhu S, Cao Z, Liang Y, Sboner A et al (2014) PRC2 is recurrently inactivated through EED or SUZ12 loss in malignant peripheral nerve sheath tumors. Nat Genet 46: 1227–1232

Li P, Silvis MR, Honaker Y, Lien WH, Arron ST, Vasioukhin V (2016) alphaE-catenin inhibits a Src-YAP1 oncogenic module that couples tyrosine kinases and the effector of Hippo signaling pathway. Genes Dev 30: 798–811

Liberzon A, Birger C, Thorvaldsdottir H, Ghandi M, Mesirov JP, Tamayo P (2015) The Molecular Signatures Database (MSigDB) hallmark gene set collection. Cell Syst 1: 417–425

Lingo JJ, Elias EC, Quelle DE (2025) Novel Therapeutics and the Path Toward Effective Immunotherapy in Malignant Peripheral Nerve Sheath Tumors. Cancers (Basel) 17: 2410

Love MI, Huber W, Anders S (2014) Moderated estimation of fold change and dispersion for RNA-seq data with DESeq2. Genome Biol 15: 550

Ma S, Meng Z, Chen R, Guan KL (2019) The Hippo Pathway: Biology and Pathophysiology. Annu Rev Biochem 88: 577–604

Macagno N, Pissaloux D, de la Fouchardiere A, Karanian M, Lantuejoul S, Galateau Salle F, Meurgey A, Chassagne-Clement C, Treilleux I, Renard C et al (2022) Wholistic approach: Transcriptomic analysis and beyond using archival material for molecular diagnosis. Genes Chromosomes Cancer 61: 382–393

Martin E, Coert JH, Flucke UE, Slooff WM, Ho VKY, van der Graaf WT, van Dalen T, van de Sande MAJ, van Houdt WJ, Grunhagen DJ et al (2020) A nationwide cohort study on treatment and survival in patients with malignant peripheral nerve sheath tumours. Eur J Cancer 124: 77–87

O’Hare T, Shakespeare WC, Zhu X, Eide CA, Rivera VM, Wang F, Adrian LT, Zhou T, Huang WS, Xu Q et al (2009) AP24534, a pan-BCR-ABL inhibitor for chronic myeloid leukemia, potently inhibits the T315I mutant and overcomes mutation-based resistance. Cancer cell 16: 401–412

Paudel SN, Hutzen B, Cripe TP (2023) The quest for effective immunotherapies against malignant peripheral nerve sheath tumors: Is there hope? Mol Ther Oncolytics 30: 227–237

Piccolo S, Panciera T, Contessotto P, Cordenonsi M (2023) YAP/TAZ as master regulators in cancer: modulation, function and therapeutic approaches. Nat Cancer 4: 9–26

Raudvere U, Kolberg L, Kuzmin I, Arak T, Adler P, Peterson H, Vilo J (2019) g:Profiler: a web server for functional enrichment analysis and conversions of gene lists (2019 update). Nucleic Acids Res 47: W191–W198

Rebe C, Ghiringhelli F (2019) STAT3, a Master Regulator of Anti-Tumor Immune Response. Cancers (Basel) 11: 1280

Rosenbluh J, Nijhawan D, Cox AG, Li X, Neal JT, Schafer EJ, Zack TI, Wang X, Tsherniak A, Schinzel AC et al (2012) beta-Catenin-driven cancers require a YAP1 transcriptional complex for survival and tumorigenesis. Cell 151: 1457–1473

Schuetze SM, Wathen JK, Lucas DR, Choy E, Samuels BL, Staddon AP, Ganjoo KN, von Mehren M, Chow WA, Loeb DM et al (2016) SARC009: Phase 2 study of dasatinib in patients with previously treated, high-grade, advanced sarcoma. Cancer 122: 868–874

Shannon P, Markiel A, Ozier O, Baliga NS, Wang JT, Ramage D, Amin N, Schwikowski B, Ideker T (2003) Cytoscape: a software environment for integrated models of biomolecular interaction networks. Genome Res 13: 2498–2504

Si Y, Ji X, Cao X, Dai X, Xu L, Zhao H, Guo X, Yan H, Zhang H, Zhu C et al (2017) Src Inhibits the Hippo Tumor Suppressor Pathway through Tyrosine Phosphorylation of Lats1. Cancer Res 77: 4868–4880

Somaiah N, Paudyal B, Winkler RE, Van Tine BA, Hirbe AC (2024) Malignant Peripheral Nerve Sheath Tumor, a Heterogeneous, Aggressive Cancer with Diverse Biomarkers and No Targeted Standard of Care: Review of the Literature and Ongoing Investigational Agents. Target Oncol 19: 665–678

Somatilaka BN, Sadek A, McKay RM, Le LQ (2022) Malignant peripheral nerve sheath tumor: models, biology, and translation. Oncogene 41: 2405–2421

Suppiah S, Mansouri S, Mamatjan Y, Liu JC, Bhunia MM, Patil V, Rath P, Mehani B, Heir P, Bunda S et al (2023) Multiplatform molecular profiling uncovers two subgroups of malignant peripheral nerve sheath tumors with distinct therapeutic vulnerabilities. Nat Commun 14: 2696

Szklarczyk D, Nastou K, Koutrouli M, Kirsch R, Mehryary F, Hachilif R, Hu D, Peluso ME, Huang Q, Fang T et al (2025) The STRING database in 2025: protein networks with directionality of regulation. Nucleic Acids Res 53: D730–D737

Velez-Reyes GL, Koes N, Ryu JH, Kaufmann G, Berner M, Weg MT, Wolf NK, Rathe SK, Ratner N, Moriarity BS et al (2021) Transposon Mutagenesis-Guided CRISPR/Cas9 Screening Strongly Implicates Dysregulation of Hippo/YAP Signaling in Malignant Peripheral Nerve Sheath Tumor Development. Cancers (Basel) 13: 1584

Voisin L, Lapouge M, Saba-El-Leil MK, Gombos M, Javary J, Trinh VQ, Meloche S (2024) Syngeneic mouse model of YES-driven metastatic and proliferative hepatocellular carcinoma. Dis Model Mech 17: dmm050553

Wu LMN, Deng Y, Wang J, Zhao C, Wang J, Rao R, Xu L, Zhou W, Choi K, Rizvi TA et al (2018) Programming of Schwann Cells by Lats1/2-TAZ/YAP Signaling Drives Malignant Peripheral Nerve Sheath Tumorigenesis. Cancer cell 33: 292–308 e297

Wu Z, Bonneil E, Belford M, Boeser C, Ruiz Cuevas MV, Lemieux S, Dunyach JJ, Thibault P (2022) Proteogenomics and Differential Ion Mobility Enable the Exploration of the Mutational Landscape in Colon Cancer Cells. Anal Chem 94: 12086–12094

Yao C, Zhou H, Dong Y, Alhaskawi A, Hasan Abdullah Ezzi S, Wang Z, Lai J, Goutham Kota V, Hasan Abdulla Hasan Abdulla M, Lu H (2023) Malignant Peripheral Nerve Sheath Tumors: Latest Concepts in Disease Pathogenesis and Clinical Management. Cancers (Basel) 15: 1077

Zheng Y, Pan D (2019) The Hippo Signaling Pathway in Development and Disease. Dev Cell 50: 264–282

